# Asciminib mitigates DNA damage stress signalling induced by cyclophosphamide in the ovary

**DOI:** 10.1101/2020.12.30.424825

**Authors:** Luca Mattiello, Giulia Pucci, Francesco Marchetti, Marc Diederich, Stefania Gonfloni

## Abstract

Cancer treatments often have adverse effects on the quality of life for young women. One of the most relevant negative impacts is the loss of fertility. Cyclophosphamide is one of the most detrimental chemotherapeutic drugs for the ovary. Cyclophosphamide may induce the destruction of dormant follicles while promoting follicle activation and growth. Herein, we demonstrate the *in vivo* protective effect of the allosteric Bcr-Abl tyrosine kinase inhibitor Asciminib on signalling pathways activated by cyclophosphamide in mouse ovaries. Besides, we provide evidence that Asciminib did not interfere with the cytotoxic effect of cyclophosphamide in MCF7 breast cancer cells. Our data indicate that concomitant administration of Asciminib mitigates the cyclophosphamide-induced ovarian reserve loss without preventing the anticancer potential of cyclophosphamide. Altogether these observations are relevant for the development of effective ferto-protective adjuvants to preserve the ovarian reserve from the damaging effect of cancer therapies.

## 1. Introduction

Chemotherapy, radiation, or combinations of them are commonly used in cancer therapy. Ovarian failure and infertility are well-known side effects of such therapies [1–3]. Cyclophosphamide (Cy) is an alkylating chemotherapeutic drug routinely used against solid and hematological malignancies [4]. Recent studies highlighted the primary target of Cy *in vivo* in murine ovaries [5–8]. Cyclophosphamide exposure induced death of growing granulosa cells and the concomitant activation of the DNA Damage Response (DDR) and AKT-FOXO3a signaling axis in the nucleus of reserve oocytes [6]. These observations supported the hypothesis that damaging stress pathways are activated in a concomitant manner both in somatic and germ cells following chemotherapy. However, the identification of sentinel molecules directly involved in communicating stress signalling remains a daunting task, as the physiological changes in the ovary reflect both direct or indirect effects of genotoxic assaults. Despite this, understanding the molecular mechanisms underlying ovarian reserve loss induced by cancer therapies remains essential for developing a more effective treatment to preserve the fertility of female patients.

## 2. Results

### 2.1 Asciminib induced a re-localization of the c-Abl tyrosine kinase to the perinuclear zone

In this study, we evaluated the effects of Cy (or of an active metabolite 4-hydroperoxy-cyclophosphamide, 4-OH-Cy) alone and in combination with Asciminib, either *in vivo* in mice or *in vitro* in a model cell line for breast cancer (MCF7). First, we validated the re-localization of c-Abl after treatment with Asciminib as previously described for the allosteric inhibitor GNF2 [9]. We monitored the c-Abl localization in a transgenic MEF line (Mouse Embryonic Fibroblast, lacking the expression of c-Abl) following transfection with a c-Abl expression vector. We evaluated the c-Abl localization by immunofluorescence (IF) assay in transfected MEF cells, treated with different c-Abl inhibitors. We tested either allosteric ligands such as GNF2, Asciminib, and one ATP-binding competitive inhibitor, Imatinib. IF assays clearly showed that Asciminib, as well as GNF2, induced a re-localization of c-Abl in the perinuclear zone as indicated by the yellow arrows (Figure 1). On the contrary, treatment with Imatinib did not cause any enrichment of c-Abl kinase in the perinuclear zone.

**Figure 1.**
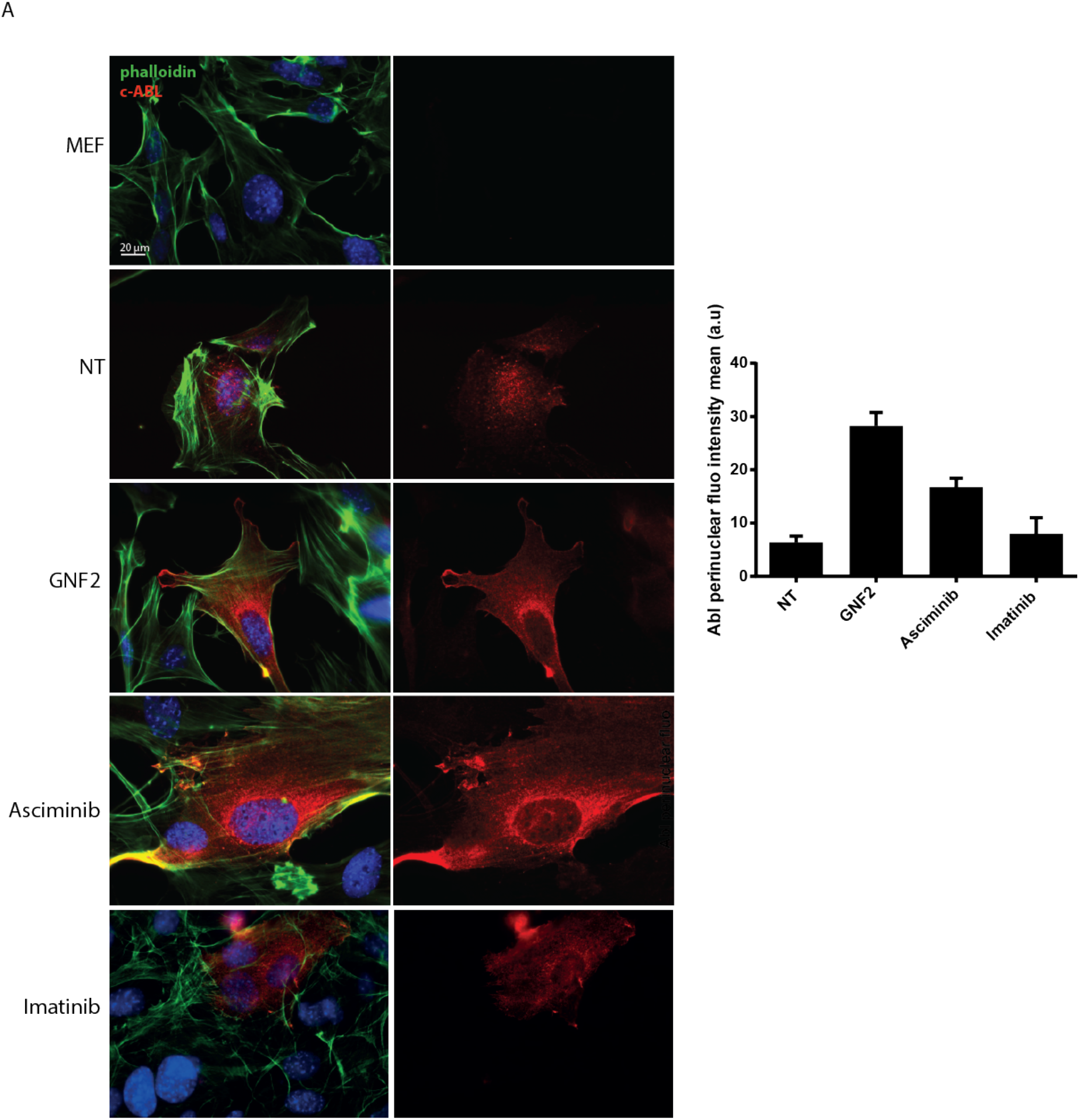
Allosteric inhibitors caused an enrichment of c-Abl tyrosine kinase in the perinuclear zone. IF assay on MEF Abl-/- cells that were transiently transfected with an expression vector for c-Abl. The yellow arrows indicated the re-localization of c-Abl tyrosine kinase in the perinuclear zone induced by Asciminib or GNF2 exposure. Bar column represents mean ± s.e.m. Scale bar 20 μm.

### Asciminib modulated the DDR and the follicle activation induced by Cy exposure *in vivo*

Next, we injected P7 mice with Cy alone (100mg/kg) or in combination with increasing concentrations of Asciminib (0.1, 0.2, and 0.5 mg/kg, respectively). At different time points, the ovaries were dissected, lysed, and analyzed by IF or western blot (W.B.) assays. Co-treatment with Asciminib resulted in partial inhibition of TAp63 phosphorylation (commonly observed as a shift of TAp63 protein by W.B. assay). In Figure 2A, we observed a partial prevention of TAp63 shift (see black arrows) 18 hours after co-injection of Cy and Asciminib. We observed the phosphorylation of histone H2AX at Ser139 (γH2AX), an early marker of DDR in the ovarian lysates. Of note, co-treatment with Asciminib affected the γH2AX phosphorylation as assessed by W.B. assay (Supplementary Figure 1).

**Figure 2.**
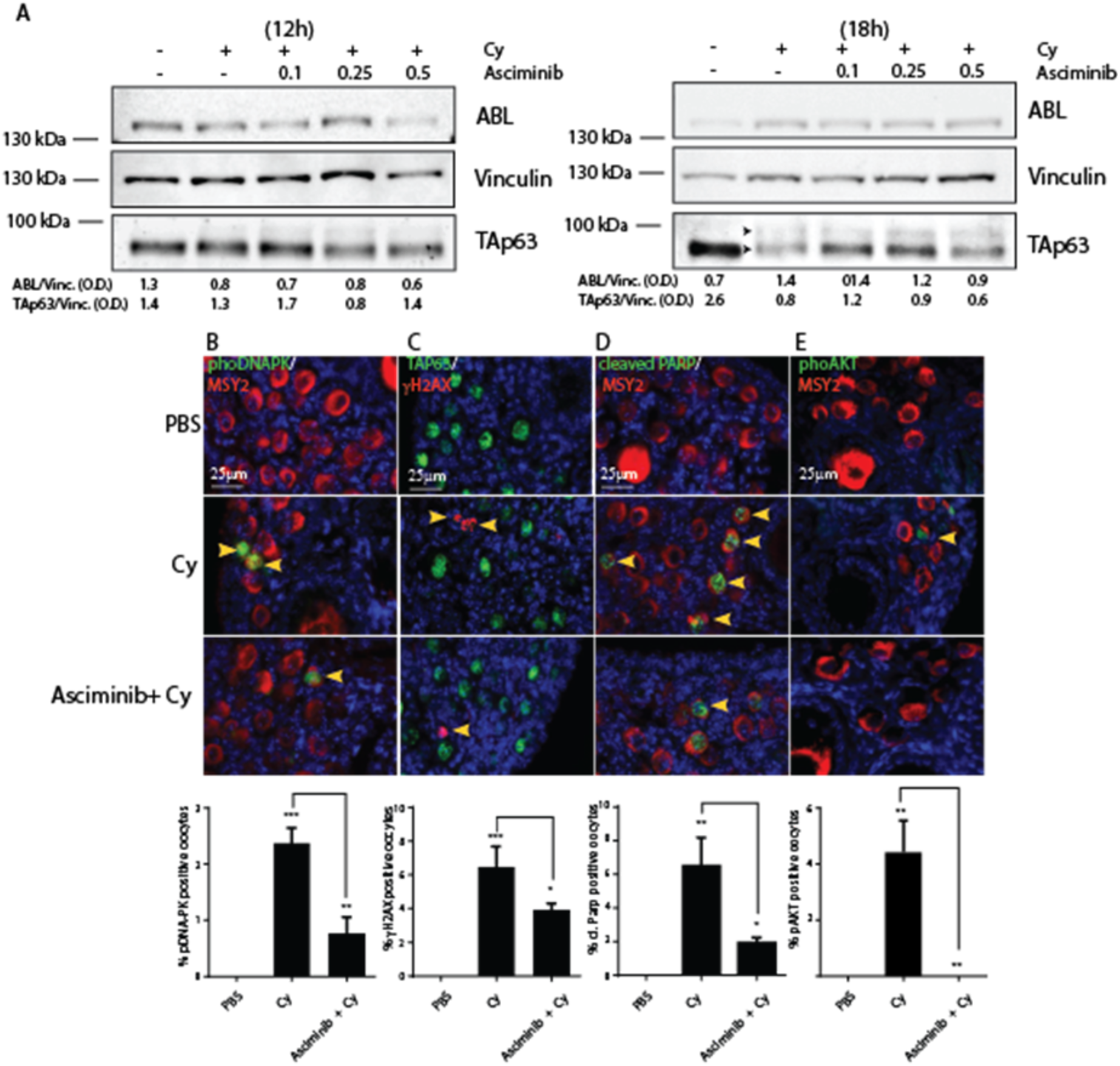
Asciminib mitigated both DDR and apoptosis induced by Cy in the ovarian reserve. (A) W.B. analysis of ovarian lysates from female pups injected with Cy 100mg/kg alone or in combination with various doses of Asciminib and collected at different time points. P7 mice were injected with vehicle (PBS) or Cy (100 mg/kg) alone or in tandem with Asciminib (0.25mg/kg) and sacrificed within 16 h from injection. Numerical values, under the blots, represent band densitometries normalized on the housekeeping gene. (B) DNA-PK activation was evaluated by IF assay using phospho-specific antibodies (green), while Msy2 (red) was used as a cytoplasmic marker for germ cells. (C) γH2AX phosphorylation was evaluated by IF assay using phosphospecific antibodies (red), while p63 (green) was used as a nuclear marker for germ cells (D) Follicle reserve apoptosis was assessed by IF assay with two specific antibodies against cleaved PARP (green) and Msy2 (red). (E) The follicular activation was assessed by IF assay with specific phospho-antibodies for AKT (T308) (green) and Msy2 (red). Quantification was performed by counting several (6 < x < 8) middle ovarian sections derived from three distinct ovaries. Scale Bar magnification, 25 μm. Bar column represents mean ± s.d.; statistical significance was determined using one-way analysis of variance (ANOVA) (*P < 0.05; **P < 0.01; ***P < 0.001 compared with PBS-treated group).

To assess whether Asciminib may affect DDR activation in primordial/primary oocytes, we monitored the phosphorylation of DDR sentinel proteins by IF assays performed on ovarian sections. We found that Asciminib attenuated DNA stress signalling induced by Cy in the ovarian reserve. Co-treated ovaries showed reduced staining for phospho-DNA-PK, γH2AX and cleaved PARP in the nucleus of reserve oocytes (Figure 2 panels B, C, D). Furthermore, co-treatment of reserve oocytes with Asciminib and Cy prevented nuclear AKT phosphorylation (Figure 2 E). Taken together, these data demonstrated that Asciminib modulated signalling pathways activated by Cy in the ovary.

### Asciminib protected the ovarian reserve following Cy treatment

Next, we injected female P6 pups with Cy (100mg/kg) alone or in combination with Asciminib (0.25mg/kg). Three days after injection, we collected ovaries to perform immunohistochemistry (IHC) assays with an antibody against cytoplasmic germ cell antigen (Msy2) (red) (Figure 3). We counted the follicle reserve from mid-ovary sections of different ovaries. IHC assays of ovarian sections showed a massive depletion of primordial and primary follicles in Cy-treated mice, whereas a Cy+Asciminib co-treatment significantly rescued reserve follicles. Follicle protection was dependent on the concentration of Asciminib, as shown in Supplementary Figure 2A, higher dosage of Asciminib (1 mg/kg) did not prevent follicle death induced by Cy and seemed to have a toxic effect *per se*.

**Figure 3.**
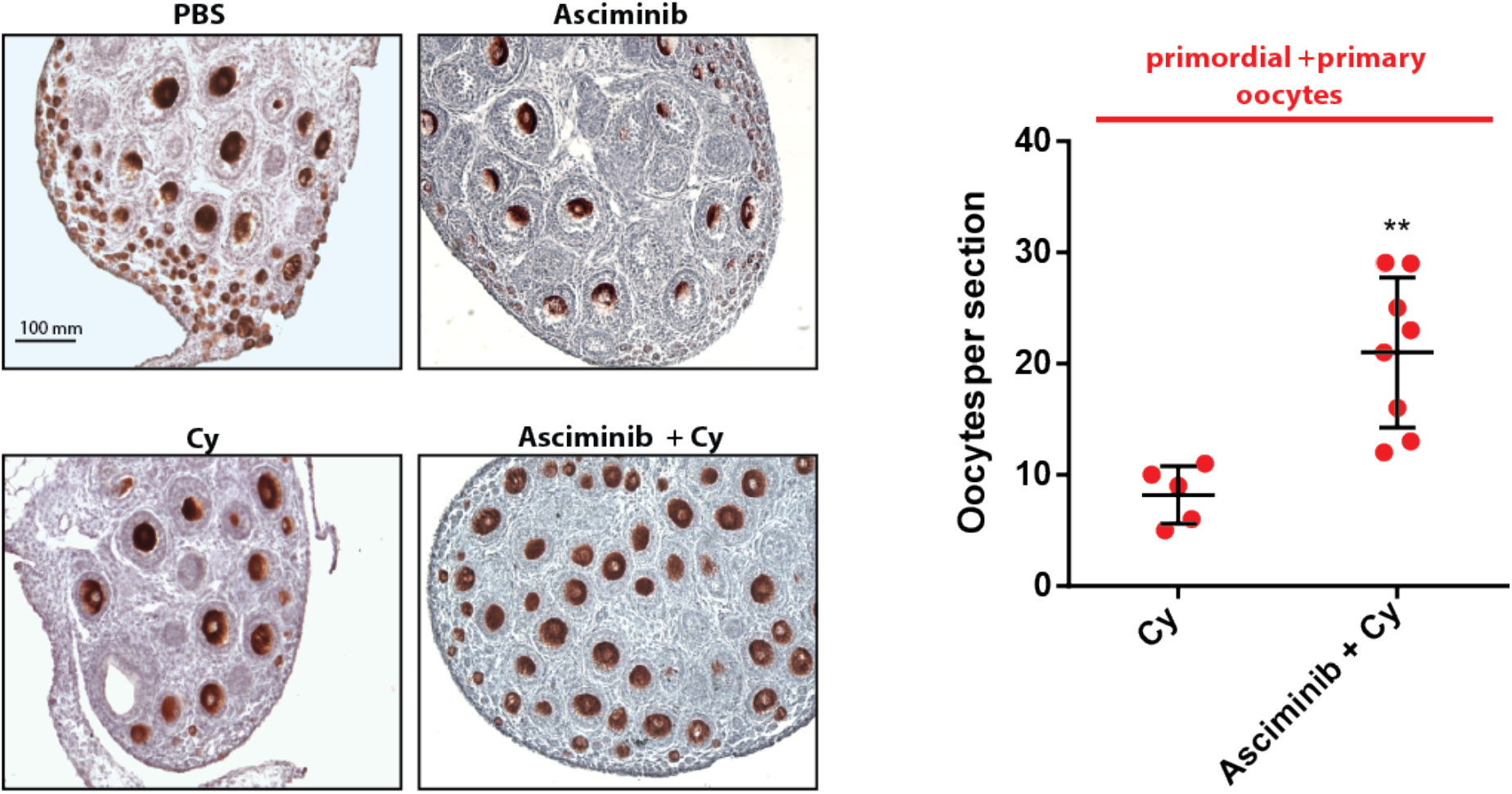
Asciminib protected the ovarian reserve from Cy treatment. Ovaries of each experimental group were dissected three days after injection (Mice were injected with Cy 100mg/kg alone or in tandem with Asciminib 0.25mg/kg) and analyzed by IHC assay with Msy2 antibody. Several ovaries from independent experiments were analyzed. Each dot in the box plot represents the average primordial+primary follicles numbers per section of each gonad collected. Statistical significance was determined using one-way analysis of variance (ANOVA) (**P<0.01 compared to Cy 100 mg/kg). Scale Bar, 100 μm.

### Asciminib did not prevent the DNA damage induced by Cy in MCF7

A clinically-used ferto-protective drug should not interfere with the therapeutic effect of DNA-damaging chemotherapies. We validated this assumption by assessing the effect of Asciminib on 4-hydroperoxy-cyclophosphamide (4-OH-Cy)-treated MCF7 breast tumor cells. Our results showed that the co-treatment with Asciminib did not affect 4-OH-Cy-induced phosphorylation of DDR marker proteins like ATM, γH2AX or p53 (Figure 4A). Besides, single-cell gel electrophoresis (Comet) assays showed that Asciminib did not interfere with the DNA-damaging effect of 4-OH-Cy (Figure 4B). Lastly, the co-administration of Asciminib did not affect the cytotoxic effect of 4-OH-Cy (Figure 4C). Altogether these data support the potential use of Asciminib as a *ferto-protective* drug without abrogating the cytotoxic effect of 4-OH-CY.

**Figure 4.**
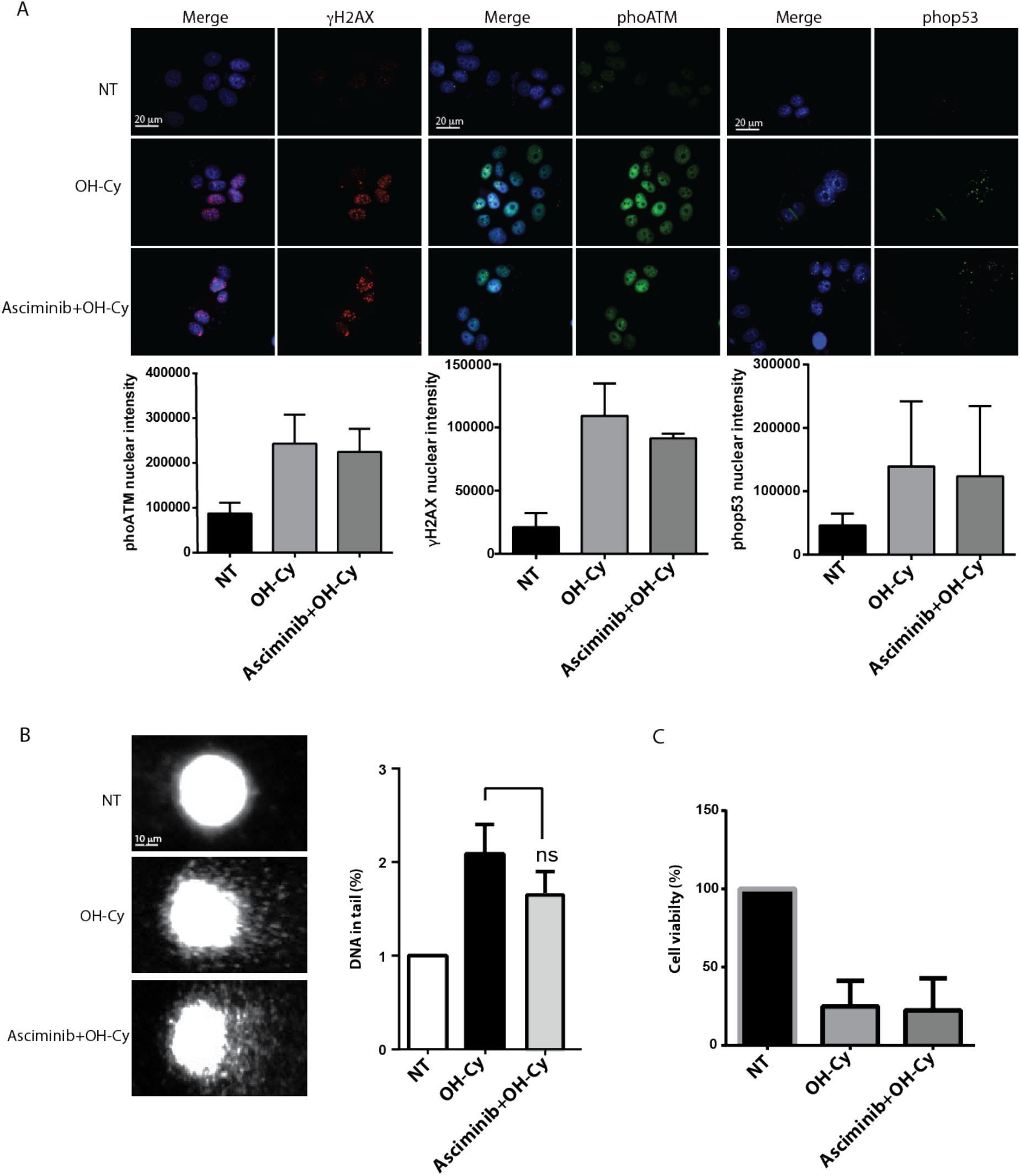
Asciminib did not prevent the cytotoxicity of 4-OH-Cy in MCF7. MCF7 cells were treated with Asciminib (0.5 μm). After 1 hour, 4-OH-Cy was added in the medium (4-OH-Cy 10 μM for comet and IF assay, 4-OH-Cy 50 μM for MTS assay. (A) DDR signalling was evaluated by IF after 4 hours of 4-OH-Cy treatment using specific phospho-antibodies for sentinel proteins. Bar columns represent mean ± s.e.m.; nuclear fluorescence intensity was evaluated by ImageJ software. Scale bar 20 μm. (B) DNA fragmentation was assessed by comet assay following 4 hours of 4-OH-Cy treatment; DNA percentage in tail quantification was evaluated by Comet Score software. Bar columns represent mean ± s.e.m.; Scale bar 10 μm. (C) Drug toxicity was measured by MTS assay after 48 hours of 4-OH-Cy treatment. Bar columns represent mean ± s.e.m.

## 3. Discussion

In young women, chemotherapy regimens increase the risk of premature ovarian failure (POF) and infertility. Genotoxic agents have two different effects on ovarian function: the first one is immediate, induces amenorrhea and the loss of growing follicles; the second one has a long-term impact by inducing a loss of the ovarian reserve. While the first effect is reversible, the loss of the follicle reserve is permanent and leads to infertility. As such, fertility preservation is an urgent issue for cancer survivors. Pharmacological agents can prevent follicle loss at the time of treatment while providing several advantages over fertility conservation techniques. These small molecules could act as ferto-protective drugs [10] by counteracting the effects of chemotherapy on the ovarian reserve. Ferto-protective drugs may be suitable for patients of all ages while avoiding the use of invasive hormonal and surgical procedures. Such ferto-adjuvants should not interfere with chemotherapeutic treatments while preventing endocrine side effects of premature ovarian failure and infertility [11]. Primordial follicles maintain their genome integrity for several decades in human without relying on classical DNA quality check controls of somatic cells during the cell cycle [12]. A better understanding of the signalling pathways activated in the follicle reserve (formed by a single oocyte surrounded by few granulosa cells) following chemotherapy is fundamental for the development of effective ferto-protective drugs. Recent findings from Rinaldi *et al* suggested that the oocytes reaching a threshold level of unrepaired DSBs after irradiation can be eliminated by a signalling pathway that required both p53 and TAp63α transcription factors [13]. Compelling evidence suggested a role of c-Abl in the oocyte degeneration induced by chemotherapy [14–16]. In mice, pharmacological inhibition of c-Abl counteracted the cytotoxic effect of cisplatin [14, 16, 17] and cyclophosphamide [6]. In this study, we tested a novel selective allosteric compound targeting c-Abl, Asciminib, against the damaging effects of cyclophosphamide. Allosteric inhibitors bind a deep pocket in the large lobe of the c-Abl catalytic domain, which is very far from the ATP binding site. Of note, small allosteric ligands induced conformational changes even in the ATP binding pocket of the kinase, preventing its active conformation [18]. Asciminib is a novel and more selective inhibitor compared to GNF2 [19]. Also, Asciminib is already used in phase III assays (NCT03106779) against resistant forms of Chronic Myeloid Leukemia (CML) [20]. Here, we showed that the co-administration of Asciminib had a protective effect on the ovarian reserve, as assessed by immunohistochemistry (IHC) performed three days after Cy injection. Besides, we found that Asciminib did not counteract the genotoxic effect exerted by an active Cy active metabolite (4-OH-Cy) on breast cancer cells. Together these data support the potential use of Asciminib as a ferto-adjuvant during chemotherapeutic regimens.

## 4. Materials and Methods

### Animals and injection

All procedures involving mice and care have been conducted at the Interdepartmental Service Centre-Station for Animal Technology (STA), University of Rome “Tor Vergata”, in accordance with the ethical standards, according to the Declaration of Helsinki, in compliance with our institutional animal care guidelines and following national and international directives (Italian Legislative Decree 26/2014, Directive 2010/62/E.U. of the European Parliament and of the Council). The ovaries were collected from CD-1 mice (Charles River) of 6 to 8 days old. Newborn mice (P6) were treated with intraperitoneal (I.P.) injection with PBS or Cy (100 mg per kg of body weight). Mice were pre-treated with Asciminib (0,1-0,5 mg per kg of body weight). Cy (BAXTER) was fresh prepared at 40 mg/ml in PBS. We dissolved Asciminib in DMSO.

### Immunohistochemistry, Follicle counting and statistical analysis

We prepared sections from ovaries fixed in MetaCarnoy solution, embedded in paraffin and cut in slices of 7 μm of thickness. Sections were dewaxed, re-hydrated, and microwaved. Slices were then permeabilised with PBS triton 0,2 % and incubated with MSY-2 antibody (Santa Cruz). The staining was performed with immunocruz staining system for anti-goat antibody (Santa Cruz, sc-2023) and 3-aminoethyl-9-ethylcarbazole as substrate (AEC, Sigma). Sections were counterstained with hematoxylin and cover-slipped with Aquatex. Quantification of primordial and primary or secondary follicles was derived from histological analysis, counting Msy2-positive germ cells of mid-ovary sections. For each ovary (derived from P9 mice P9 pups), several central slices (10<n<15) are included in the counting, except for smaller peripheral slices (12-14 on average per each ovary). Quantification of primordial/primary follicle reserve is expressed as mean of immature follicles (primordial plus primary follicles) per single ovary. Average values for each ovary are represented as discrete points on a scatter plot. Mean value ± S.D. are shown in the scatter plot. The analysis of variance is evaluated with one-way ANOVA, with Turkey multiple comparison Test using PRISM 6 (Graph Pad software) (*P<0.05; **P<0.01; ***P<0.001) or by unpaired Student’s t test where indicated.

### Immunofluorescence

We prepared sections from ovaries sections fixed in MetaCarnoy solution, embedded in paraffin and cut in slice of 7 μm of thickness. Sections were dewaxed re-hydrated and microwaved in sodium citrate 10mM pH6, to expose the antigens. Unspecific-binding sites were blocked by incubating sections for 2 hours in a blocking solution (PBS plus 1% glycine 5% FBS and 5% NGS (normal goat serum). Ovaries sections were then incubated overnight with antibodies against MSY-2, p63, γH2AX, p-DNA-PK, pAKT and cleaved PARP. After washing in PBS triton 0,05%, tissue sections were incubated with Alexa 555-goat anti-mouse (life technologies) and Alexa 488-goat anti rabbit (Invitrogen). Nuclei were stained with 1 μg/mL Hoechst 33342 dyes for cells (Thermo Fischer Scientific) in PBS 1X.

### Immunoblot analysis

P7 dry ice-frozen ovaries were homogenized with a mini-pestle in ice-cold lysis buffer (50 mM Tris-HCl pH 7.5, 150 mM NaCl, 0.5% NP-40, 5 mM EDTA, 0.5% sodium deoxycholate, 1 mM phenylmethylsulfonyl fluoride, 1 mM sodium o-vanadate, 10 μg ml-1 Tosyl phenylalanyl chloromethyl ketone (TPCK), 10 μg ml-1, Tosyl-L-lysyl-chloromethane hydrochloride (TLCK) supplemented with protease inhibitors, (all purchased from SIGMA). Equal amounts of protein extract (equivalent of one up to three ovaries) was loaded onto 6%, 8% or 12% SDS-PAGE gel and transferred to a nitrocellulose membrane (Amersham Bioscience). Immunoblot densitometry were performed using ImageJ software.

### Cell culture

MEF Abl-/- cells were kindly gifted by the Koleske lab (Yale, USA). MCF7 cells were kindly provided by the Barilà group (IRCCS-Fondazione Santa Lucia, Rome, Italy). Cells were grown in DMEM medium (GIBCO) supplemented with 15% FBS (Mef Abl-/-) or 10% FBS (for MCF7) (Lonza) and 100 U/ml penicillin/streptomycin (Lonza) in a humidified atmosphere containing 5% CO_2_ at 37°C.

### Cells transfection and Immunofluorescence assay

MEF Abl-/- were grown in a 6-wells plate on cover slips and transfected with a plasmid encoding wild-type c-Abl 1b isoform for 16 hours (at low passages), by using lipofectamine 2000 DNA Transfection Reagent (Invitrogen) according to the manufacturer’s instruction. During this time the cells were also treated with small-molecule c-Abl compounds alone (DPH, GNF-2, Asciminib) for 4 hours and then analyzed by Immunofluorescence.

MCF-7 cultured in a 6-wells plate on cover slips were incubated for 60 minutes with allosteric c-Abl inhibitors, Asciminib (0.5 μM) or GNF-2 (10 μM), treated with 4-hydroperoxycyclophosphamide (4-OH-Cy) (sc-206885) for 4 hours and analyzed by Immunofluorescence.

Cells were grown on cover slips, then washed in PBS and fixed with 4% paraformaldehyde for 10 min at room temperature. Cells were incubated with a solution containing Triton X-100 (0.5%), blocked for 2 hours with a blocking solution (PBS, Triton X-100 0.1%, BSA 5%), and then incubated with primary antibodies against p-ATM, γH2AX and p-p53 for 60 minutes. After washing in PBS/Triton 0,1%, cells were incubated with Alexa 555-goat anti-mouse (Life technologies) and Alexa 488-goat anti rabbit (Invitrogen) for 30 minutes. Nuclei were stained with 1 μg/ml 4,6-diamidino-2-phenylindole dihydrochloride (Thermo Fischer Scientific). Fluorescence images were obtained by Leica DMR Fluorescence Microscope (Leica Microsystems, Germany). Perinuclear fluorescence analysis was performed using FIJI software. In particular, the regions of interest (ROI) were identified using DAPI signal to spot the edge of nuclear region. From this edge, a 30 pixels-wide area was pointed out using the fix ellipse command and then used as ROI to measure fluorescence intensity mean.

### Single cell gel electrophoresis assay (Comet Assay)

Breast cancer cells (MCF7) cultured in 60 mm dishes were incubated for 60 minutes with allosteric inhibitors, Asciminib (0.5μM) or GNF-2 (10μM), treated with 4-hydroperoxycyclophosphamide (OH-Cy) for 4 hours. All following steps were conducted under dim light to prevent the occurrence of additional DNA damage. The cells were washed with PBS twice and 20μl of each cellular lysate were mixed with 730μl of 0,5% low melting agarose solution. One tenth of this volume was dropped on slides coated with 1% normal melting agarose. The slides were covered with coverslips and placed on ice for 5 min. After gel solidification coverslips were gently removed. The slides were placed into Schifferdecker type glass cuvette, filled with lysis solution (10 mmol/L Tris-HCl, 2.5 mol/L NaCl, 100 mmol/L EDTA, 1% Triton-X 100, 10% DMSO, pH 10, 4°C) and incubated at 4°C overnight. After lysis step, the slides were washed with deionized water and placed into electrophoresis chamber (Apelex) filled with 2.2 L of alkaline electrophoretic solution (300 mmol/L NaOH, 1 mmol/L EDTA, 4°C, pH>13,) for 30 min. Electrophoresis was performed in the same solution for 20 min at electric field strength of 0,7 V/cm. The applied voltage was 11 V and the current was 280 mA. After electrophoresis, the slides were washed twice with neutralization buffer (Tris-HCl 0,4M pH 7.5), fixed in 70% ethanol for 10 minutes, dried at room temperature and stored until staining. Immediately prior to microscopic analysis, the slides were stained with GelRed (SIGMA) in the dark. The images of comets were analyzed using Comet Score software and approximately 300 cells per slide were counted. DNA fragmentation was evaluated by the percentage of DNA in the tail of comet (% DNA tail).

### MTS Assay

MCF7 cells were seeded into 96-wells plates at a density of 10000 cells per well (200 μl) and treated with the following: vehicle control (DMSO), OH-Cy alone or in combination with Asciminib (0.5 μM). The cells were treated for 24 h. For the MTS assay, the CellTiter 96® AQueous One Solution Cell Proliferation Assay kit (Promega, USA) was used following the manufacturer’s instruction. The absorbance of cells for each condition was detected at 490 nm with a microplate reader (Sunrise Tecan microplate reader).

### Reagents

Antibodies for Msy-2 (sc-21316) and p-AKT (T308) (sc-16646-R) were purchased from Santa Cruz; antibody for p-H2AX (γH2AX) (05-636), H2AX (07-627) and were purchased from Millipore; antibody for p-DNA-PK (S2056) (SAB4504169) was purchased from SIGMA, antibody for p-ATM (S1981) (200-301-500) was purchased from Rockland, polyclonal antibody for p63 was a homemade rabbit serum; antibody for p-P53 (S15) (9284) and cleaved PARP (9544) were purchased from Cell Signalling Technology. Western Blot secondary antibodies were purchased from Jackson Immunoresearch. All the antibodies were diluted in a blocking solution containing 5% BSA in PBS tween 0,05% for Western Blotting analysis and in a blocking solution containing 1% glycine, 5% FBS and 5% NGS for immunofluorescence assay.

## Author Contributions

L.M. performed the majority of the experiments with the help of F.M. G.P. performed MTS and IF assays in MCF7 for phoATM, phop53, γH2AX. M.D. provided reagents and critical reading of the paper. S.G. designed, directed the study and wrote the paper.

## Acknowledgments

We thank Luisa Castagnoli for her valuable support. We are indebted to Stefano Cannata e Sergio Bernardini for technical advice. M.D. is supported by the National Research Foundation (NRF) (grant number 019R1A2C1009231), by a grant from the MEST of Korea for Tumour Microenvironment Global Core Research Center (GCRC) (grant number NRF-2011-0030001), by the Creative-Pioneering Researchers Program through Seoul National University (Funding number: 370C-20160062), by the Brain Korea 21 (BK21) PLUS programme, by the ‘Recherche Cancer et Sang’ foundation, by the ‘Recherches Scientifiques Luxembourg’ association, by the ‘Een Haërz fir kriibskrank Kanner’ association, by the Action LIONS ‘Vaincre le Cancer’ association and by Télévie Luxembourg.

